# A model-based analysis of the mechanical cost of walking on uneven terrain

**DOI:** 10.1101/2020.06.15.152330

**Authors:** Alexandra S. Voloshina, Arthur D. Kuo, Daniel P. Ferris, C. David Remy

## Abstract

Human walking on uneven terrain is energetically more expensive than on flat, even ground. This is in part due to increases in, and redistribution of positive work among lower limb joints. To improve understanding of the mechanical adaptations, we performed analytical and computational analyses of simple mechanical models walking over uneven terrain comprised of alternating up and down steps of equal height. We simulated dynamic walking models using trailing leg push-off and/or hip torque to power gait, and quantified the compensatory work costs vs. terrain height. We also examined the effect of swing leg dynamics by including and excluding them from the model. We found that greater work, increasing approximately quadratically with uneven terrain height variations, was necessary to maintain a prescribed average forward speed. Greatest economy was achieved by modulating precisely-timed push-offs for each step height. Least economy was achieved with hip power, which did not require as precise timing. This compares well with observations of humans on uneven terrain, showing similar near-normal push-off but with more variable step timing, and considerably more hip work. These analyses suggest how mechanical work and timing could be adjusted to compensate for real world environments.

## Introduction

It is energetically more expensive for humans to walk on natural, complex terrain than on hard, flat surfaces [1–5]. A multitude of factors, such as terrain unevenness, damping, stiffness, friction, and other surface characteristics can affect gait biomechanics and energy expenditure. Such potential causes for increases in energy consumption can be roughly classified into two categories: gait adaptations that facilitate control or preempt falls when walking under more challenging conditions and gait adaptations that are mechanically necessary to account for terrain variation. The first category includes, for example, increased muscle co-activation for greater joint stiffness during walking on slippery surfaces [6–8], or a more crouched posture for improved stability during running [9]. Such adaptations might lead to sub-optimal gait through a voluntarily made trade-off between gait stability and gait economy. In contrast, increases in energetic cost associated with mechanically necessary adaptations on uneven terrain cannot be offset by alternative gait adjustments. Even when walking at the theoretical optimum, surface properties can require more mechanical work to be performed against the ground. For example, walking on energy-dissipating surfaces, such as sand, necessitates more mechanical work than walking on hard surfaces [10]. Similarly, surface height variations impact center of mass trajectories [11], which may lead to greater energy expenditure [12]. It is often difficult to experimentally distinguish the specific contribution of a gait adaptation to an increase in energetic cost, since humans likely make multiple adaptations at the same time. An alternative way to investigate how uneven terrain affects gait energetics is through a model-based analysis that focuses on one single gait characteristic.

To illustrate the effects of center of mass motion on energy expenditure, consider a simple example of moving along an upward step of height *d* followed by an equal downward step of height −*d*. When the center of mass moves upwards, potential energy changes by Δ*E* = *mgd* (with *m* being the total body mass and *g* being the gravitational acceleration). Without any other adaptations, this change in potential energy must be generated by mechanical work, *W*, performed over the course of the step, such that *W* = Δ*E* = *mgd*. As the downwards step is associated with an equal amount of negative work −*W*, the net energetic effect of the two steps is zero. There may, however, be metabolic energy expended for negative work, if actively performed by muscle. It has been estimated that when humans walk up and down steep inclines they perform positive work with an efficiency of *η*^+^ = 25 % and negative work with an efficiency of *η^−^* = −120 % [13]. That is, generating positive or negative mechanical work can require positive net metabolic effort. The amount of additional metabolic energy *E*_VCOM_ to move up and down vertically would be:

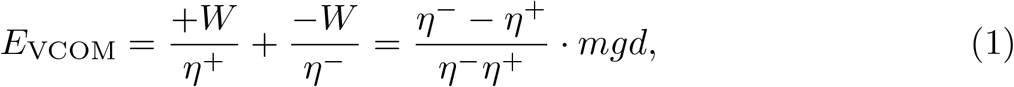

and has a net positive magnitude. However, this does not account for the possibility of gait adaptations that might occur in response to the irregular surface, and an optimal gait that mitigates some of the effects of Equation (1) likely exists. For example, strategies other than direct work production could account for fluctuations in potential energy. By slowing down when going up a step and speeding up when going down, potential energy could be gained from kinetic energy, in turn reducing the amount of necessary active work [14]. Similarly, taking advantage of collisions to passively perform negative work could further reduce the necessary metabolic effort. In addition to influencing the mechanical work that must be performed, such adaptations also affect average locomotion speed. To fully understand these dynamics and to identify an optimal walking strategy over irregular terrain requires a detailed model-based analysis.

The amount of work necessary for steady-state locomotion of a model on uneven terrain strongly depends on how the gait is powered. Past research has shown that energy expenditure of simulated locomotion is sensitive to the relative timing of active push-off done by the trailing limb and collision during heel-strike of the leading limb [15]. Work production is most effective when done through push-off immediately before collision of the leading leg, otherwise additional work must be done at the hip [16]. On uneven terrain, unperceived irregularities may disrupt the relative timing of push-off and heel-strike and demand work production at the hip. In addition, and especially on highly irregular terrain, theoretically optimal push-off magnitudes may actually be biologically unfeasible; although push-off and collision work must always be positive and negative, respectively, in biological gait, theoretically optimal gaits may dictate otherwise. This means that there exists a limit to the maximum amount of positive work that can be done through push-off, after which additional work must be done through the hip.

In this study, we isolated the effects of surface unevenness on energy expenditure by studying walking of simulated legged models over terrain of increasing surface variability. Specifically, we evaluated the energetic consequences of the mechanical adaptations required to move up and down over irregular terrain, and the mechanisms by which mechanical work can be performed to do so. To this end, we investigated two simple models of legged locomotion walking over uneven surfaces: the powered rimless wheel [17] and the powered simplest walking model [18–20]. For simplicity, we represented uneven terrain as a surface consisting of alternating up and down steps of equal height. Both models were powered by impulsive push-off and hip work. Similar models have been able to closely approximate, among other parameters, center of mass and joint dynamics, ground reaction forces, and swing leg motion during locomotion on level ground [17, 21, 22]. In addition, such models have made suggestions for economical methods to power locomotion [14, 15]. For the two models, we analytically calculated the effects of terrain irregularity on step timing and average forward speed, and numerically simulated a range of adaptation strategies. We then identified optimal locomotion patterns and calculated the effective costs of transport assuming different combinations of push-off and hip work to actuate gait. Finally, we noted if increases in cost of transport (COT) were similar between model and empirical human data, allowing us to evaluate if humans are able to locomote in an optimal fashion over irregular terrain. Together, these analyses provide a clearer understanding of the bipedal adaptation methods and their energetic consequences on uneven terrain.

### Description of Mechanical Models

We adapted two simple inverted pendulum models of walking to investigate how cost of transport is affected by varying terrain variability, gait types, and methods of energy input. The powered rimless wheel [17] takes steps of fixed length and has no swing leg dynamics. It is modeled as rigid spokes fixed at an inter-leg angle of 2*α* (Fig. 1A). In contrast, the powered simplest walking model [19, 20] includes passive swing leg dynamics, and is modeled as two rigid legs connected at the hip by a frictionless hinge joint (Fig. 1A). Both models have legs of length *l*, a point mass *m* located at the hip (or wheel center) and representing the pelvis and torso, and feet with negligible mass *m_f_*. Stance leg dynamics for both models are simply that of an inverted pendulum:

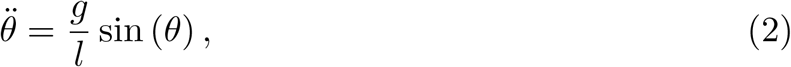

where *θ*(*t*) is the stance leg angle relative to vertical and *g* is gravity.

**Figure 1.**
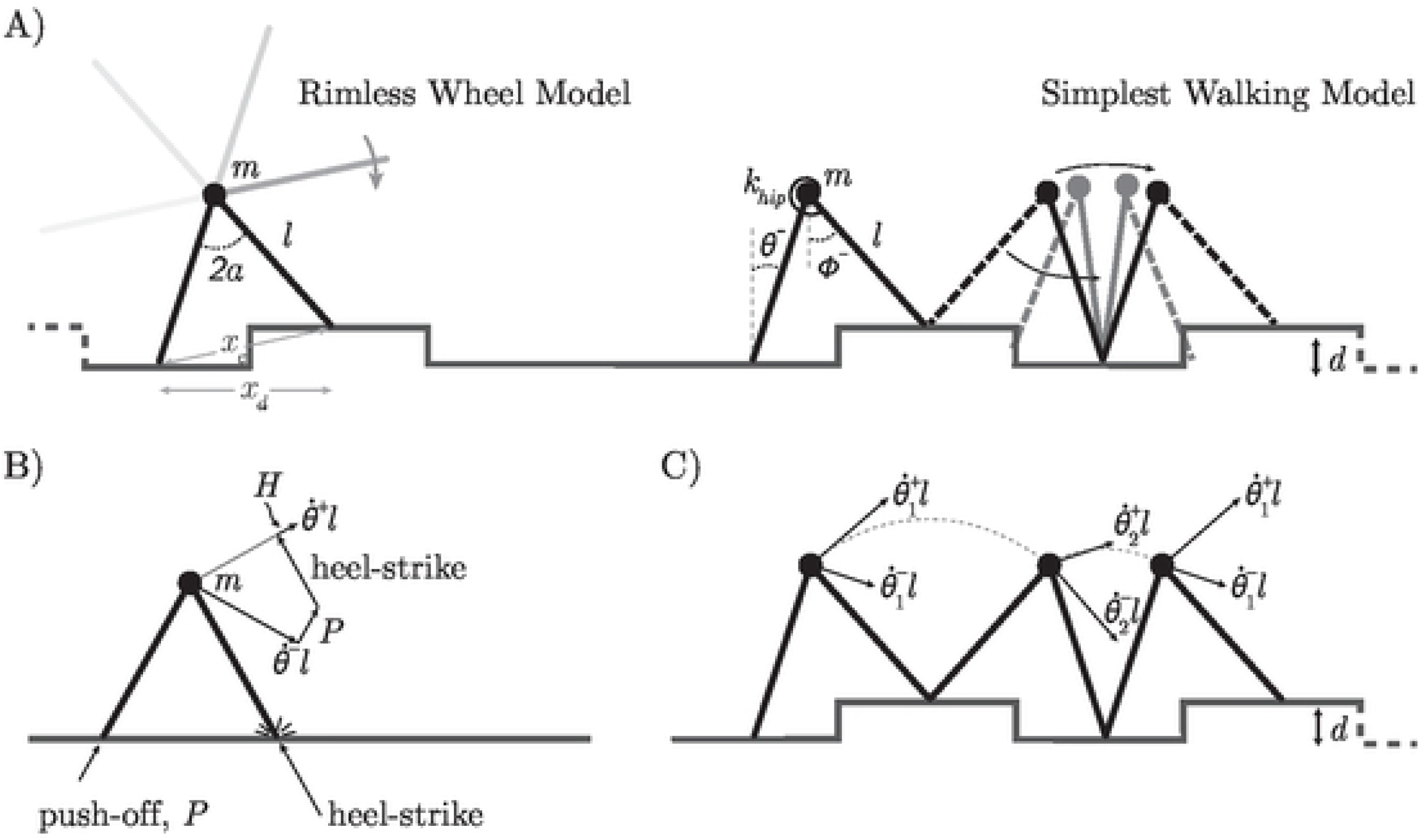
Schematic of the powered rimless wheel and simplest walking models. A) Models have a hip mass *m*, massless legs of length *l* and feet of negligible mass *m*_*f*_ (not shown). Uneven terrain consists of up and down steps of height *d*, with *x*_*e*_ and *x*_*d*_ defined as step lengths on even and uneven ground, respectively Rimless wheel spokes are 2*α* degrees apart. Simplest walking model swing leg dynamics are determined by the torsional hip spring *k*_hip_ B) Energy input occurs through a push-off impulse *P,* hip impulse *H,* or both Before collision, the center of mass (COM) moves with velocity 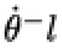. Push-off *P* is applied along the trailing leg, redirecting the COM velocity. The heel-strike impulse then further redirects COM velocity to be perpendicular to the new stance leg. Impulsive hip work *H,* directed along the motion of the COM after collision, brings the COM velocity after collision to the desired initial velocity of the step, 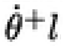. C) Diagram of velocity indexing for the up and down steps.

For the simplest walking model, *ϕ*(*t*) is the swing leg angle relative to vertical. A torsional hip spring (with stiffness *k_hip_*) applies a torque between the stance and swing legs at a desired swing leg frequency, 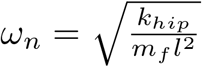. The torsional spring is analogous to hip flexor and extensor muscles activity observed during steady human walking. Assuming that *m_f_ ≪ m*, the swing leg does not affect stance leg dynamics [15], and swing leg dynamics for the simplest walking model reduce to:

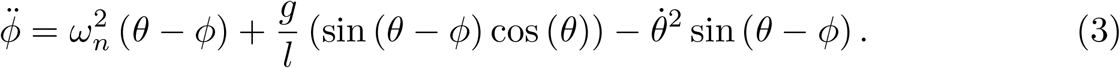

When the leading leg contacts the ground, the roles of the stance and swing switch, and the models undergo an instantaneous and perfectly inelastic collision that redirects the center of mass velocity to be tangential to the new stance leg. At this point, the leg angles update such that *θ*^+^ = *ϕ^−^* and *ϕ*^+^ = *θ^−^* for the simplest walking model, and *θ*^+^ = *θ^−^* + 2*α* for the rimless wheel. In these and the following equations, the superscripts ‘−’ and ‘+’ refer to states immediately before and after collision, respectively. Energy is lost due to the negative work done by the collision and must be replaced through positive work to maintain steady state walking.

Two strategies have been proposed as analogs of human lower limbs doing positive work on the center of mass: ankle push-off initiated by the trailing leg shortly before heel-strike of the leading leg, and the application of hip torques throughout the stance phase [15, 17]. In our models, we approximated both strategies using impulses (Fig. 1B). The ankle push-off impulse, 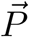, is directed along the trailing leg towards the COM and applied immediately before heel-strike of the leading leg. That is, push-off is *perpendicular* to the direction of motion of the COM *before* collision. In contrast, the hip impulse 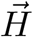 is applied *along* the COM direction of motion *after* collision. The push-off impulse acts analogously to work done by the lower leg muscles before toe-off, while the hip impulse is analogous to the hip torque produced by hip extensor muscles in the human, applied between the torso (not included in our simplified models) and stance leg [15, 23, 24].

Considering a passive collision, as well as the magnitudes of active push-off and hip impulses, post-collision velocities of the stance and swing legs can be derived with impulse-momentum. For the simplest walking model they are given by:

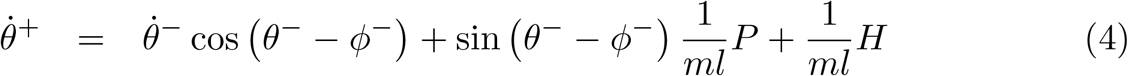

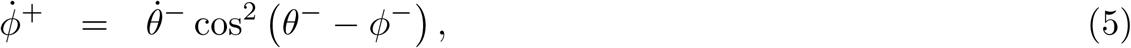

and for the rimless wheel by:

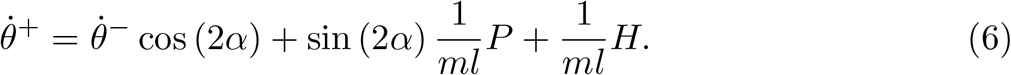

The above equations are a modification of the derivations presented in [15] and [19], and include the effect of 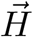 on stance leg angular velocity, also referred to as “late push-off” [14, 25].

The post-collision stance leg angular velocity, 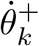, of each step *k* is thus a direct function of the pre-collision velocity, 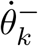, and the magnitudes of the impulses *H_k_* and *P_k_* at this step. The pre-collision velocity, 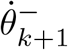, at the *subsequent* heel-strike is then obtained by integrating the continuous dynamics given by Equations (2) and (3), starting from 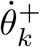 For the rimless wheel, the fixed leg angle and repeating structure of the terrain result in the same COM height at the start and end of each step, independent of whether the model is moving up or down. Since stance phase of the models is energetically conservative, the integration leads to a trivial result for the rimless wheel:

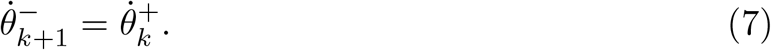

In contrast, the inter-leg angle of the simplest walking model can vary from step to step and the above relationship no longer holds. The velocity 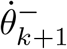 and leg angles 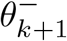 and 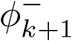 of the simplest walking model can thus only be computed via numerical integration of the equations of motion.

Given the periodic nature of the simulated surface, we restricted our analysis to two-step periodic motion and assumed that leg angles and velocities for each second step are identical, such that 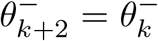 and 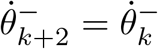. Similarly, 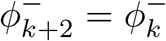 and 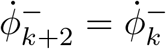 for the simplest walking model. With this, we introduce the following naming convention for stance leg angular velocities (Fig. 1C):
 
- 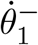 and 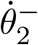: velocities immediately before collision, following a down or up step, respectively;
- 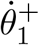 and 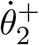: velocities immediately after collision, leading into an up or down step, respectively.

Through numerical integration, we can determine the step time, *t_k_*, and the step distance, *x_k_*, of each step by taken by the simplest walking and rimless wheel models. From this, the average speed 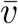 over two steps is computed as:

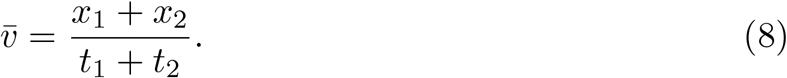

Step durations and distances are dependent on step height *d* and on the initial angular, or step, velocities 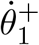 and 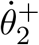. Each initial step velocity needs to be sufficiently large in order to overcome the apex of the COM trajectory and to complete the step in a finite amount of time. Without adaptions in the initial step velocities, walking over uneven terrain leads to larger average step durations, 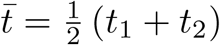, and shorter effective step lengths, 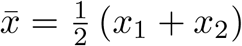. This, in turn, influences the average forward speed 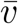 (for details, see Appendix S1). In order to understand the mechanical adaptations necessary for walking on uneven terrain, we inverted this dependency; we constrained 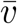 to a fixed value, analogous to walking with a constant speed on a treadmill. This means that if step height *d* is increased, initial step velocities must be adapted to maintain a desired 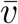. This essentially introduces an implicit relationship between the initial step velocities of the up and down steps. If the desired walking speed and one initial step velocity is prescribed, the second initial step velocity can be uniquely determined. Once we determined the relationship between 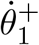 and 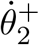 given a desired forward speed, we computed the magnitudes of *P* and *H* that achieved this relationship.

### Computation of Work and Energy

To compute the required mechanical work and associated metabolic effort to maintain a desired forward speed 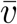, we need to quantify the work done in Equations (4) and (6). In general, mechanical work done by an impulse, 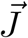, on a particle with mass *m* moving at an initial velocity 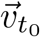, can be computed as 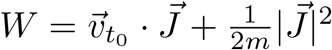. Based on the direction of push-off 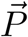 (perpendicular to the direction of motion of the COM) and hip impulse 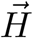 (along the direction of motion of the COM) the work of these impulses can be computed from their magnitudes according to:

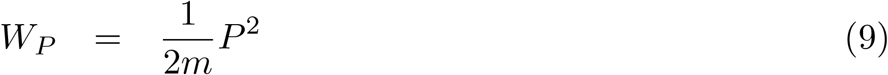

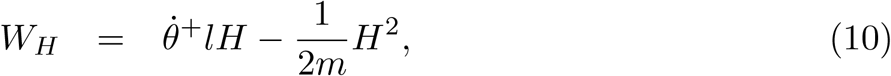

where *W*_*P*_ and *W*_*H*_ are the work done by the push-off and hip impulses, respectively (similar to [15]). The work performed by the impulsive push-off is strictly positive, while hip work can be positive or negative. If positive and negative work efficiency is defined as *η*^+^ and *η^−^*, respectively, the metabolic cost of transport (COT, non-dimensional) over two steps is given by:

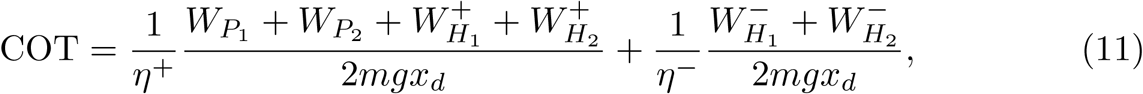

where 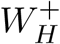 and 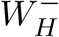 are the positive and negative hip work, respectively.

### Work Input Strategies

Gaits can be obtained through combinations of push-off and hip work that create desired post impact velocities 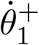 and 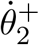 according to Equations (4) & (6). The chosen combination of impulse magnitudes *P* and *H* directly affect COT, as defined by Equation (11). To determine the effects of energy input approaches on overall energy expenditure, we evaluated rimless wheel and simplest walking model gaits powered by three different *work input strategies* that represent various abilities of anticipating the upcoming step:

1. Variable Push-off with hip work as needed (VP): Optimally timed push-off of *varying* magnitude is performed, up to the maximum possible without an aerial or jumping phase onto the next step, with hip work added beyond that point. This method likely represents a best case scenario for energy economy and presumes that all steps are fully anticipated and push-off timing and magnitude are continuously adapted. To compute the cost of transport, we first obtained the push-off magnitudes *P* that produce the required 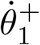 and 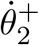 for *H* = 0. To ensure that the push-off and collision impulses had a physically realizable positive magnitude, we constrained push-off magnitude to 0 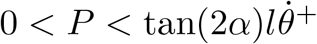, and then re-computed *H* as required.
2. Constant Push-off with hip work as needed (CP): Optimally timed push-off of *constant* magnitude is performed, again with hip work compensating to avoid aerial or jumping phase. This strategy assumes that heel-strike timing can be anticipated correctly, while step height is not adjusted for. For all step heights and velocity combinations, we set the push-off impulse magnitude to 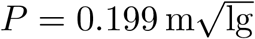, the push-off magnitude used during symmetric, level ground walking. The hip impulse then provides any additional work to achieve the required values for 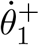 and 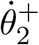.
3. Hip Only work (HO): Hip work done by hip impulse of magnitude *H*, while *P* = 0. This method presumes zero knowledge of the subsequent step and that any preemptive push-off is impossible [14].

### Computational Analysis

The rimless wheel and simplest walker models simulated gait analogous to human walking on level ground at a speed of 1.00 m/s and with a step length of 0.662 m [5]. The half-inter-leg angle (*α* = 0.390 rad) of the rimless wheel determined the step length of the model. Hip stiffness dictates step length in the simplest walking model, and was selected such that rimless wheel and simplest walking model gaits were equivalent on level ground (i.e. on level ground, 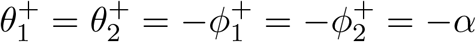). To obtain generalized results, we non-dimensionalized all outcome variables with respect to body mass (*m*), leg length (*l*, nominally 0.87 m for typical human) and gravity (*g*) as base units [26]. Hence, all gaits had an average speed of 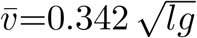, with swing leg natural frequency of the simplest walking model equal to 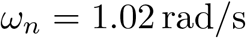.

Based on the desired gait characteristics, we numerically identified repetitive, steady-state gaits for each model that resulted in identical system states every two steps. To this end, we performed simulations over two steps, consisting of one up and one down step of equal height *d*. Numerical integration began immediately after heel-strike leading into the up step and ended immediately after heel-strike following the down step. Integration was interrupted at each heel-strike in order to define a new stance leg. At this instance, for the simplest walking model, we computed the post-collision swing foot velocity 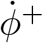 according to Equation (5). Simulations were conducted over a range of initial step velocities of the first step 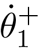, with the initial step velocity of the second step 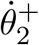 calculated such that the average forward speed over the two steps was equal to 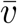. This relationship between the two initial step velocities was determined for step heights ranging from *d*=0 *l* to *d*=0.1 *l*. The simulated two-step gaits relied on a combination of push-off and hip work for power, given that any desired post-collision stance leg velocity can be achieved with an appropriate choice of impulse magnitudes *P* and *H* (per Equations (4) and (6)). Impulses *P* and *H* were restricted to follow the energy input strategies described previously. We identified all two-step periodic gaits numerically, using a nonlinear root search (similar to [27]).

## Results

We show that, given a desired average forward speed and step height *d*, the initial up and down step velocities, 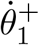 and 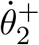, were interdependent for the rimless wheel and simplest walking models (Fig. 2). This dependency defined a range of possible gaits, which was particularly limited for the simplest walking model due to step timing constraints introduced by swing leg dynamics. For both models, cost of transport was directly related to step height and the initial up-step velocity. The relationship between these three variables was further influenced by the work input strategy (Fig. 3). In other words, the amount of total positive and negative work needed to maintain constant walking speed significantly varied if work input occurred through push-off, hip work, or a combination of the two (Figs. 4 and 5). Assuming the most economical gaits for both models and work input strategies, the rate of increase in cost of transport in response to larger step heights was larger for the simplest walking model than for the rimless wheel (Fig. 6). Finally, we compare these model results to observations of humans walking on uneven terrain [5].

**Figure 2.**
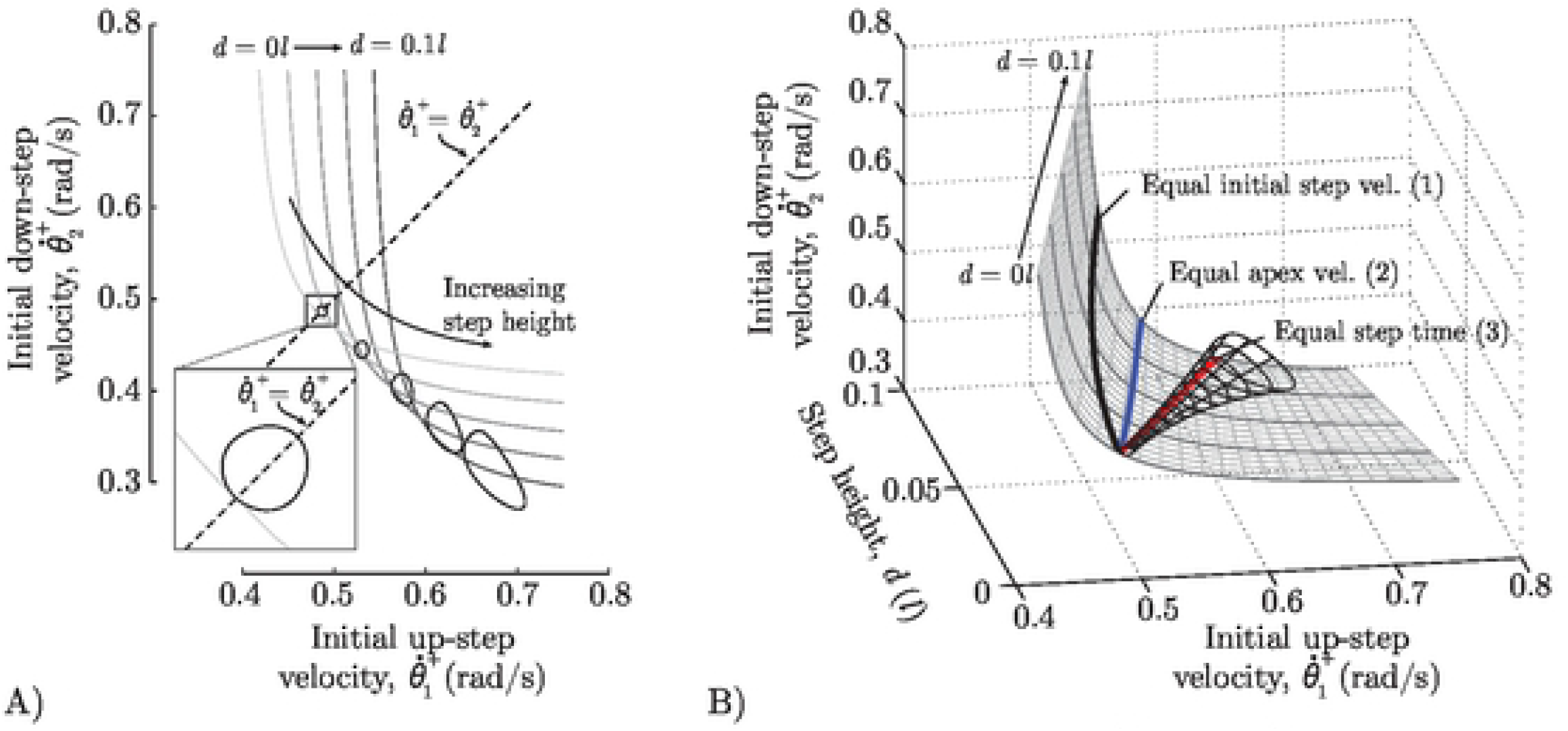
**Relationship between initial step velocities 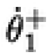 and 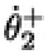, and changes in step height, *d,* for the rimless wheel and simplest walking models walking at an average forward speed of 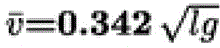.** A) Lines depict 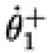 and 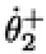 relationships for the rimless wheel (linear subspace) and simplest walking models (circular subspace) at selected step heights *d*, ranging from 0 *l* to 0.1 *l* at equal intervals of 0.025 *l*. B) The same relationship shown in a 3-dimensional space. Thicker lines on the surface correspond to the lines plotted in A). Colored lines highlight gait characteristics for particular choices of 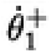 and 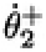.

**Figure 3.**
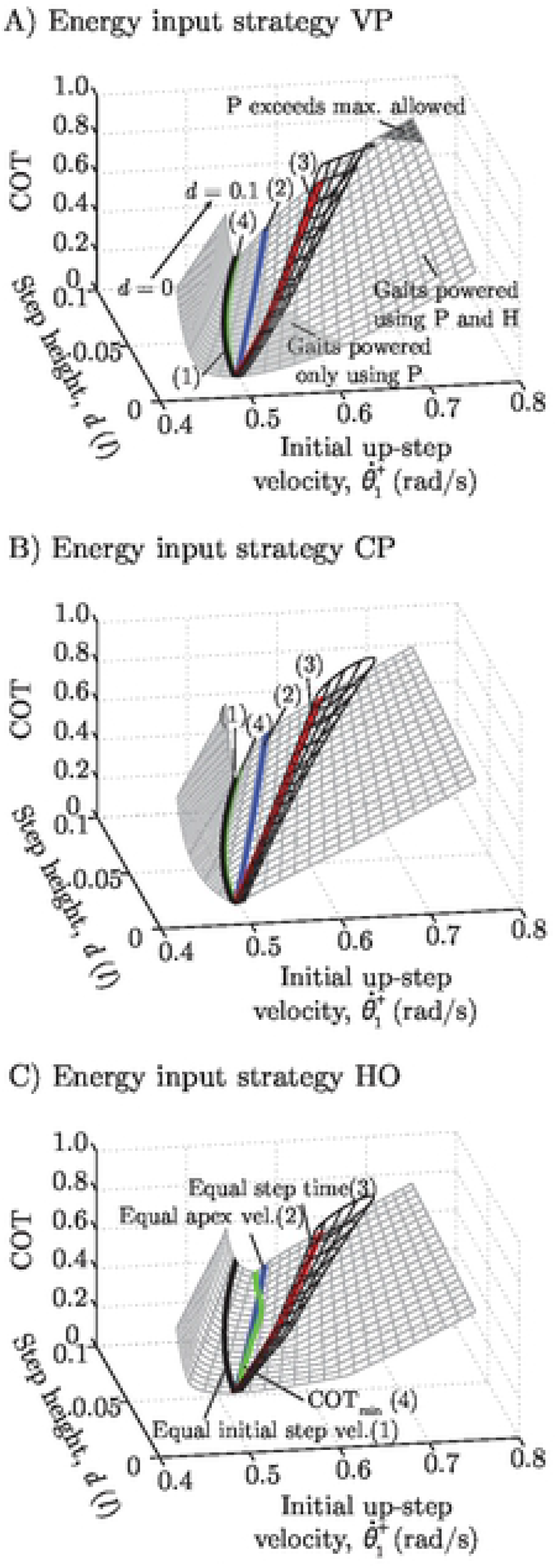
Effects of step height and work input strategy on metabolic cost of transport (COT) of the rimless wheel and simplest walking models. The COT is shown as a function of step height and the initial step velocity for the up-step. COT values are non-dimensional. The initial velocity for the down-step follows from the constraint that the average locomotion speed 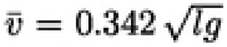. Rimless wheel solutions lie on a planar manifold while simplest walking model solutions lie on a tubular manifold. Compared are three work input strategies: A) Variable Push-off with hip work as needed (VP), B) Constant Push-off with hip work as needed (CP), and C) Hip Only work (HO). Solid green lines mark the energetically optimal choice of 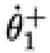 and 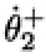 for the rimless wheel model, *COT*_min_. The remaining colored lines highlight gait characteristics for particular choices of 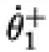 and 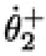 (that result in gaits with equal initial step velocities (1), equal velocities at the top of each step (2), equal step durations (3), and minimum energetic cost).

**Figure 4.**
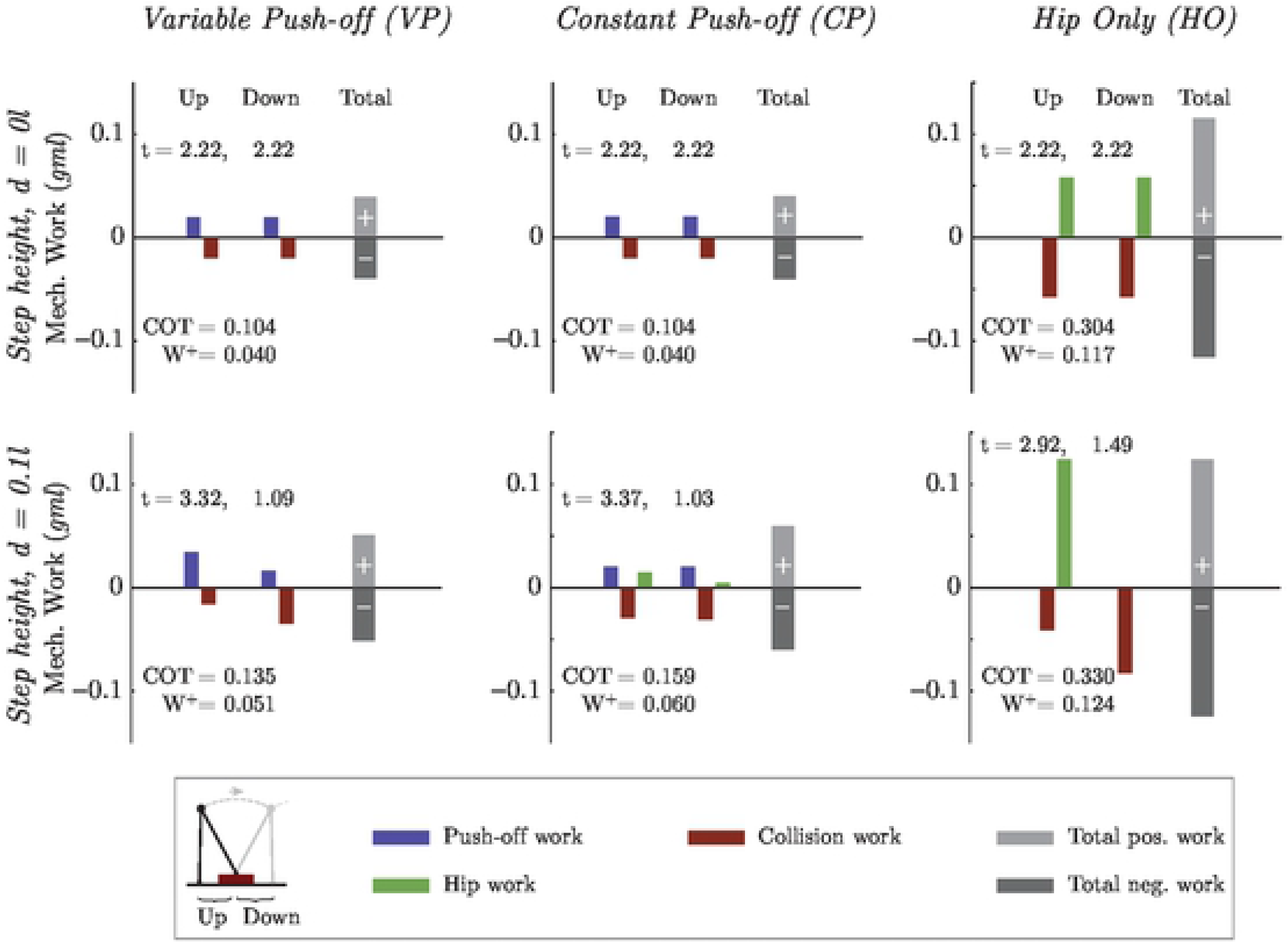
Breakdown of work contributions for the rimless wheel during up and down steps, given varying work input strategies on even and uneven terrain. Work breakdowns are shown for energetically optimal solutions (green line for each respective work input strategy (Fig. 3)), for up and down step velocities, 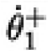 and 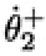. Average walking speed over two steps is 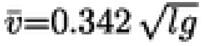. Cost of transport (COT) and work (*W*^+^) values are non-dimensional. Note that no active negative work is required during optimal rimless wheel gaits. Step durations are non-dimensionalized with respect to gravity and leg length (expressed in units 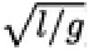).

**Figure 5.**
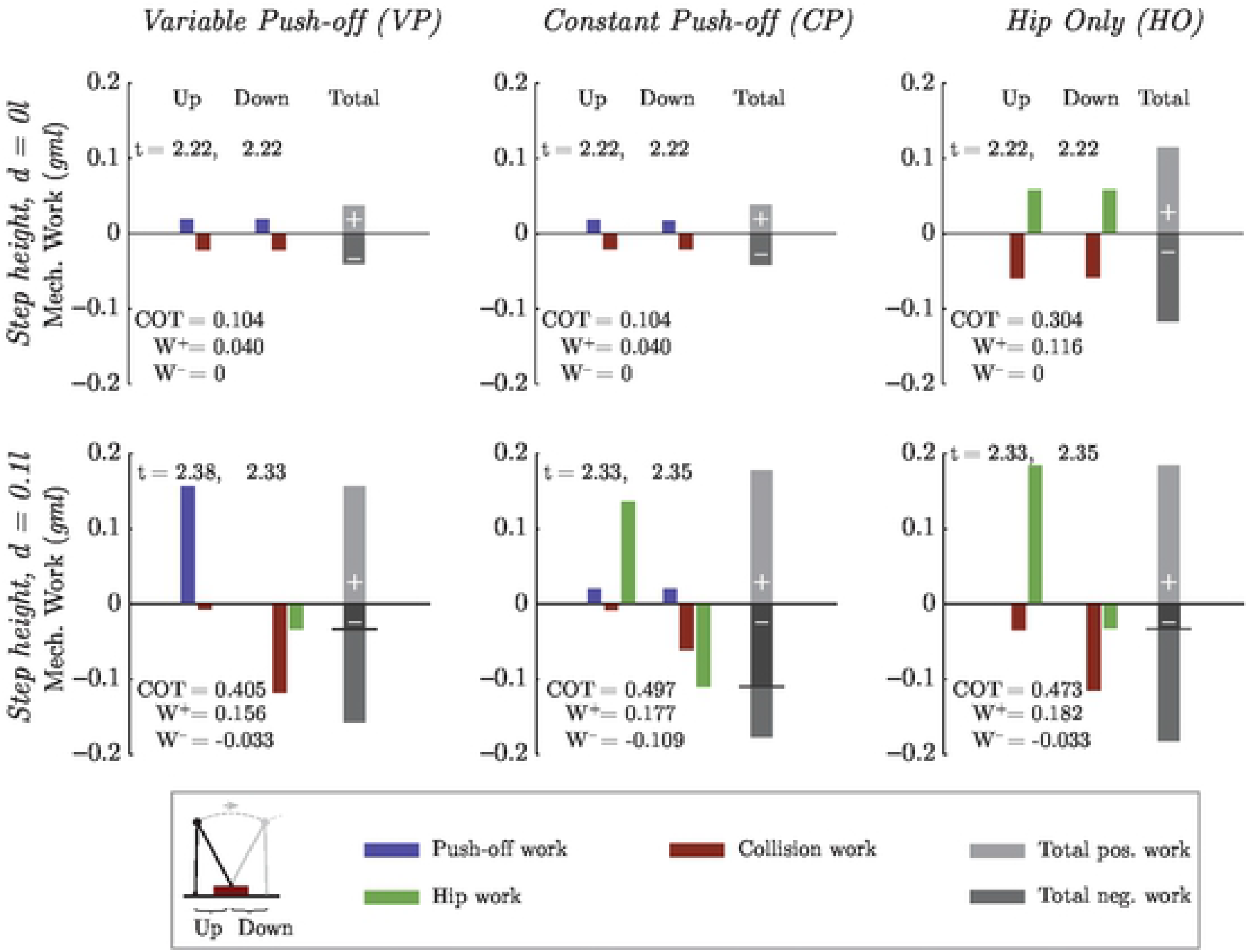
Breakdown of work contributions for the simplest walking model during up and down steps, given varying work input strategies on even and uneven terrain. Work breakdowns are shown for energetically optimal solutions (green line for each respective work input strategy (Fig 3)), for up and down step velocities, 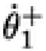 and 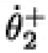. Average walking speed over two steps is 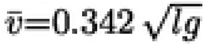. Cost of transport (COT) and work (*W*^+^ and *W*^−^) values are non-dimensional. Note that, unlike the rimless wheel, simplest walking model optimal gaits require active negative work to be performed during walking on uneven terrain. Step durations are non-dnnensionalized with respect to gravity and leg length (expressed in units of 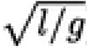). Dark shaded regions of total negative work bars indicate negative work actively performed by the hip, which contributes to total COT.

**Figure 6.**
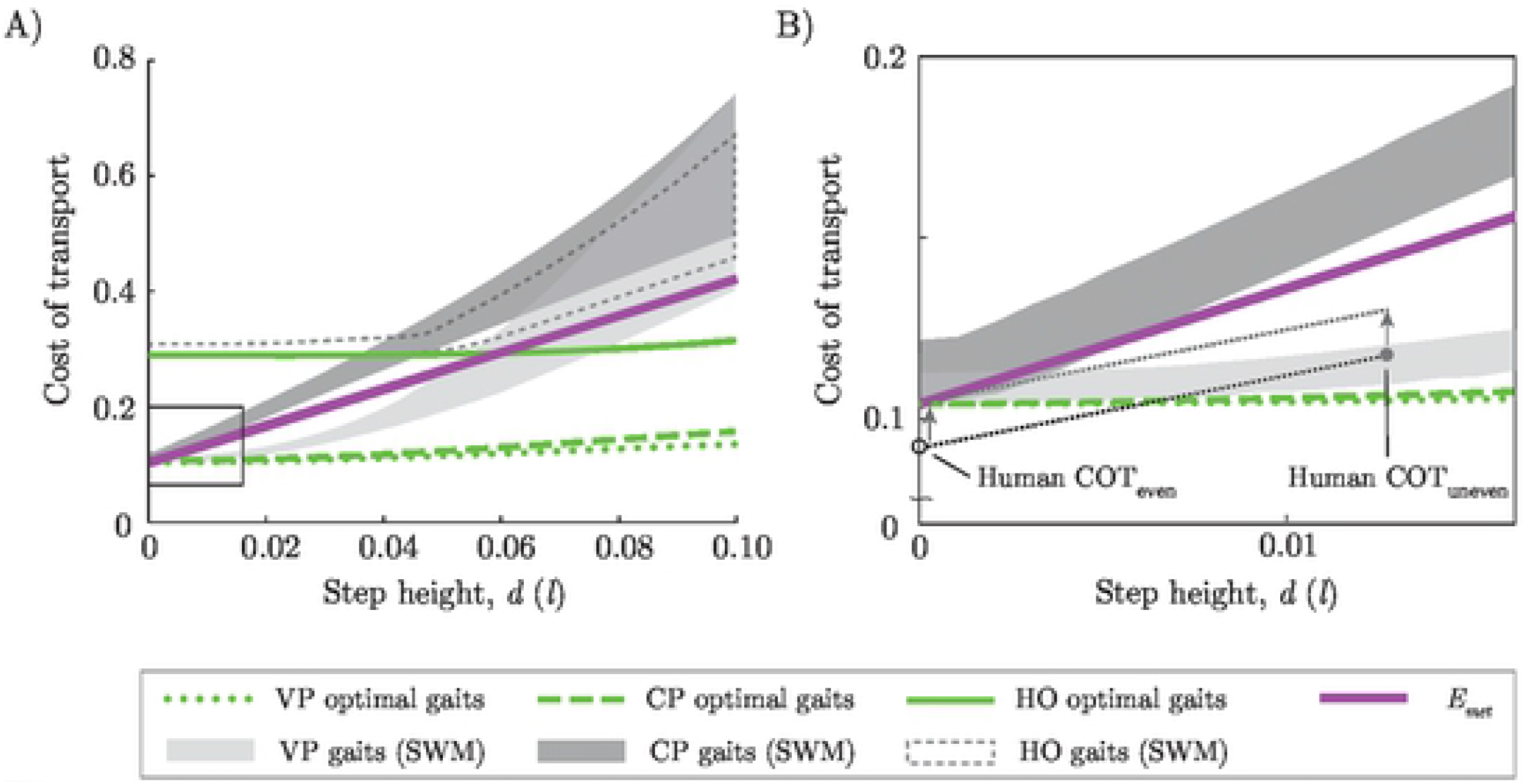
Comparison of model predicted increases in cost of transport with changes in step height. **A)** Energetically optimal gaits for rimless wheel (lines) and simplest walking model (shaded areas for all gaits, optimal gait at lower bound). Expected energy cost *E*_VCOM_ (purple line), for moving the center of mass vertically up and down the step (Equation 1). **B)** Close-up of area identified by the rectangle in **(A)**, including human metabolic cost of transport on level and uneven terrains (hollow and solid dots, respectively).

### Rimless Wheel

All possible gait solutions for the rimless wheel model lay on an extended surface in the space defined by 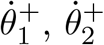, and step height *d* (Fig. 2). To maintain the desired average speed of 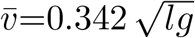, smaller values in 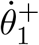 required larger values in 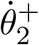 and vice versa. On level ground, the relationship between the initial step velocities was symmetric about the point where the two velocities were equal (Fig. 2A, insert). Increasing *d* shifts the symmetry point in the direction of a larger initial up-step velocities and a smaller down-step velocity.

The most energetically economical choice of initial step velocities 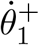 and 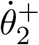 depended on the work input strategy. When powered by positive variable (VP) or constant (CP) push-off impulses, the most energetically optimal solution was practically indistinguishable from gaits with identical initial step velocities, independent of step height (Fig. 3A,B). In contrast, for gait powered only through hip work (HO), the most energetically optimal gaits were those with equal apex velocities at mid-step (Fig. 3C). These gaits generally required a larger initial velocity at the up-step than on the down step. Gaits with with equal step durations were most energetically costly, independent of work input strategy. The VP work input strategy tended to be the most economical overall, but only allowed for a limited region of gaits to be powered purely by push-off (light grey shaded part of the surface in Fig. 3A). As initial step velocities 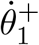 and 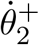 diverged, one of the push-off impulses grew, while the other got smaller. Since push-off impulses can only be positive, the smaller impulse had to be replaced by negative work done at the hip as soon as it reached a value of zero (unshaded part of the surface). Furthermore, for highly unbalanced steps 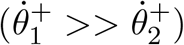 and large values of *d*, the push-off prior to the up-step became so large that it required a negative collision impulse for the rimless wheel to remain on the ground. In this region (dark grey shaded part of the surface) push-off values had to be constrained and, in order to maintain the required average forward speed, supplemented with positive hip work.

Differences in energy expenditure between the three work input strategies may be examined in terms of the positive and negative work done at every step (Fig. 4). For the energetically optimal VP-powered gait (green line, Fig. 3), larger step heights *d* required increasing push-off before the up-step and decreasing push-off before the down-step. Since a push-off impulse helps reduce impact velocity, larger push-offs were accompanied by smaller subsequent collision losses. Conversely, smaller push-offs resulted in larger energetic losses during the subsequent collision. This imbalance in push-off impulse magnitude led to an increase in total positive work from 0.040 *gml* at *d*=0 *l* to 0.051 *gml* at *d*=0.1 *l*. Gaits powered with variable push-off had the lowest COT.

On level ground, CP- and VP-powered gaits have the same push-off values and, hence, the same work distribution. As *d* grew, additional hip work was required to supplement the constant push-off of CP-powered gaits, specifically for up-steps. Hip work was always positive, and it resulted in larger collisions than VP-powered models. As a result, total positive work generated by CP-powered gaits (0.060 *gml* at *d*=0.1 *l*) was higher than the work produced by VP-powered gaits during walking on uneven terrain.

In contrast to CP- and VP-powered gaits, HO-powered gaits can only provide work input after collision, leading to much larger energy losses during collision. As *d* increased, more hip-work was done in the up-step and less in the down-step, with no hip work done in the down-step at very high values of step height. As a result, HO-powered gaits required more positive work for walking on level (0.116 *gml* at *d*=0 *l*) and uneven (0.124 *gml* at *d*=0.1 *l*) ground when compared to gaits powered by other work input strategies. This is consistent with past analyses of the effects of using hip work on total energy expenditure [20]. On level ground, powering gait only through hip work led to an almost three-fold increase in COT compared to other work input strategies. However, the rate of growth in total positive work as a function of *d* was smaller for HO-powered gaits compared to the VP- and CP-powered gaits. None of the energetically optimal rimless wheel gaits required negative hip-work, independent of work input strategy. Thus, optimal COT results are equivalent to the positive work performed.

All of the strategies exhibited increasing COT with step height, although varying considerably in magnitude (Fig. 6). The rimless wheel with variable push-off had least COT, but even constant push-off was more economical than the simplest walking model. In addition, the COT increased approximately quadratically with step height (*R*^2^ > 0.99) for all rimless wheel models, including with costlier HO-power.

### Simplest Walking Model

In contrast to the rimless wheel, the simplest walking model did not allow for gaits with significant differences in up and down step duration. This is because step timing was dependent on time required for the passive swing leg to travel forward. Consequently, simplest walking model solutions had a smaller range of possible initial step velocities 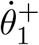 and 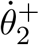. In fact, the range of possible velocities formed a closed loop in the 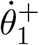 and 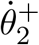 space (Fig. 2A, insert). Over increasing step height *d*, the loop widened to allow for gaits with more variable step durations (Fig. 2B).

Similarly, simplest walking model COT surfaces also formed a tubular manifold, which lay in proximity to rimless wheel gaits with equal step durations (Fig. 3). As a result, the COT of the energetically optimal gaits of simplest walking model was significantly higher compared to optimal gaits of the rimless wheel, independent of work input strategy. Simplest walking model gaits with the largest difference in step time tended to be the most energetically optimal on surfaces with larger step heights *d*. Similar to the rimless wheel, optimal VP-powered gaits were most economical for the simplest walking model at all step heights when compared to gaits using other work input strategies (Fig. 6, lower bound of shaded regions). However, in contrast to the rimless wheel, optimal HO-powered gaits had lower COT values than CP-powered gaits at higher values of step height *d* (all *d* greater than ≈0.05*l*).

On level ground, the most energetically optimal simplest walking model gaits for each work input strategy required the same amount of positive work as their respective rimless wheel counterparts (Fig. 5, first row). As step height increased, the simplest walking model required more positive and negative work during the up and down steps, respectively, when compared to the rimless wheel. This was caused by the need to maintain similar step durations, which consequently led to larger total negative work requirements. At lower step heights, and for VP- and HO-powered gaits, the increase in negative work could largely be attributed to greater, passive collision costs. At higher step heights, much of the negative was also actively generated by the hip. This is in contrast to the rimless wheel, for which negative work was always passive and did not contribute to the total COT. For CP-powered gaits, negative work was done actively through the hip even at lower step heights. This led to CP-powered gaits being more energetically expensive than HO-powered gaits at larger step heights (Fig. 5, second row), since additional negative hip work required by CP-powered gaits was more costly than the additional positive work required by HO-powered gaits.

## Discussion and Conclusions

We explored the mechanical effects of terrain unevenness on the cost of transport for the rimless wheel and simplest walking models. We simulated an uneven surface of equal up and down steps, and found periodic gaits for both models over a range of step heights. Uneven terrain could entail greater mechanical work for three main reasons. First, collision losses and cost of transport increase approximately quadratically with step height variations, for fixed step lengths. Those losses must then be restored through positive work, most economically with push-off and less so with hip work. Second, uneven surfaces limit the ability to predict heel-strike events, because push-off is most economical if timed pre-emptively before heel-strike [15, 16]. Otherwise, less economical hip work (or late push-off, [14, 25]) would be needed to maintain nominal speed. Third, swing leg dynamics also influence collision losses. The passive swing leg of the simplest walking model yields steps of varying length and duration, in contrast with the fixed step lengths of the rimless wheel. More variable steps have higher collision losses on average, and therefore require more positive work. It might be less costly overall to actively control the swing leg, depending on the trade-off its energetic cost [28] against a passive swing leg.

Although uneven terrain is more costly, it is fundamentally similar to level ground. In an inverted pendulum model, kinetic and potential energy are exchanged conservatively, and positive work adds to that energy. The only dissipation is in the heel-strike collision. This remains the case on uneven terrain, except for greater collision losses. Down-steps dissipate more energy, and up-steps less energy than nominal gait. Up-steps lose more time and speed, and the opposite for down-steps. But the differences are asymmetric, so that the losses are greater, and require more positive work with step height, even if there is no net height change over the two steps.

It is instructive to compare our model results with observations of humans walking on uneven terrain. The most relevant human experiments include mechanical and metabolic human data on level ground and on uneven terrain, with a 2.54 cm maximum step height [5], roughly analogous to the present model with *d* = 0.013 l. The net metabolic rate of walking humans increased by approximately 28% (from 2.65 W/kg to 3.38 W/kg, respectively) [5]. This corresponds to a COT increase from 0.093 to 0.118 (Fig. 6).

Such comparison allows for quick elimination of two models: vertical center of mass motion and hip-powered gait. For comparable height variation, humans walk with considerably less cost than the vertical center of mass model (*E*_VCOM_ in Equation (1) and Fig. 6), and the HO-powered strategy. The *E*_VCOM_ cost ignores all joint motion or forward motion of the body, and yet still predicts higher costs than human. Other models show uneven terrain may be negotiated for less cost than the vertical motion would imply ([14, 25]), and previous studies show that *E*_VCOM_ is a poor predictor for even level walking [12]. The HO-powered gait is far more costly than humans, even on level ground (Fig. 6). This is supported by both models and empirical data, which show that it is disadvantageous to input work via hip alone [15, 16, 29, 30]. We therefore do not consider either model explain the metabolic cost of human walking on level or uneven terrain.

The remaining models all suggest that humans should use push-off to power gait. The models gain better economy by applying pre-emptive push-off, for both level and uneven terrain. And if push-off cannot be timed precisely and varied appropriately, it is still more economical to apply constant push-off with added hip work (CP-powered gait), rather than hip work alone. On comparison terrain (*d* = 0.013 l), humans produce ankle push-off power comparable to level ground, albeit with more variable timing (about 25% more step period variability, no significant differences in ankle work [5]). It is unclear whether human push-off is limited by ankle power or poorer timing, but the additional work needed for uneven terrain appears to be supplied by the hip (about 60% more positive hip work [5]). None of the models make good numerical predictions for human costs, but they do provide a mechanistic basis for using push-off, and they approximately predict how costs should increase with step height (Fig. 6).

The models also suggest that it would be uneconomical for humans to allow the swing leg to move passively. The resulting collision losses are considerably higher than with fixed step lengths (VP simplest walking model vs. rimless wheel, Fig. 6). The human swing leg does have dynamics, unlike the rimless wheel, but can also be actively controlled, unlike the simplest walking model. Humans appear to actively control the swing leg and step length [31], with a metabolic cost [28]. That cost might be preferable to the collision losses predicted for a passive swing leg.

A limitation of this study was the use of a regular pattern of equal up and down steps for uneven terrain. It would be more realistic to model stochastic terrain structure. This would be expected to have little effect on the examination of work input strategies, since the strategies only assumed either full or zero ability to adapt to the terrain. In addition, simulations with stochastic terrains show very similar trends to the patterned terrain, albeit with greater effects on speed and other variables such as step duration or COT (see Appendix S1). Another limitation is that we also modeled relatively small terrain variations, which would be expected to induce less timing variability than than larger variations. For greater step heights, we would expect push-off timing to be more critical, and more likely to require hip work. As an extreme case, a single very high step or sequence of up-steps could bring the model to a complete halt, and require additional work input. We suspect larger step heights might cause hip work to become more dominant.

As a further limitation, we excluded the question of control from our study, focusing only on periodicity and not stability. In particular, our models relied on simple impulse control. However, simple model analysis shows an increased risk of falling on stochastic terrain [32]. Control optimization methods for walking models have been proposed to improve gait stability on uneven terrain [33, 34], and push-off could be optimized to simultaneously maintain stable gait and minimum work input [14, 25]. More sophisticated control methods could be used to study trade-offs between energy economy and perturbation rejection.

Our models are also drastically simplified compared to human. Notably, humans perform more positive and especially negative work at the knee on uneven terrain [5], and may dissipate collision losses more actively than modeled here. It would also be helpful to model the torso, which can be useful for stabilizing walking in response to perturbations [35]. We also assumed a fixed relationship between work and energy expenditure, which is quite complex in human. Not only are there non-work contributions to energetic cost such as force production ([28]) and muscle co-activation, but other factors such as body posture, altered gait kinematics (e.g. to increase swing foot clearance), and soft tissue deformation [36] can all affect energy expenditure. It is possible that performing an energetic analysis with a more complex model and the addition of gait stability analysis and control, would allow more insight into how humans balance the trade-off between stability and energetic cost.

Uneven terrain poses increasing mechanical demands on bipedal walking. There appear to be multiple ways to meet those demands by performing work to maintain forward speed despite unevenness. In our models, trailing leg push-off appears to be an economical means to add work, but only if it can be timed appropriately, and not limited by saturation or other constraints. Hip work is less economical, but also far more forgiving of timing. Humans appear to retain similar push-off on uneven terrain, but more of the modulation appears to take place with hip. Simple models cannot capture the complexity and multiple degrees of freedom of the human body, but they may provide basic insights regarding how to walk under challenging conditions.

## Acknowledgments

This work was supported by the National Science Foundation, Alexandria, VA, USA, under Grant No. CMMI 1453346 to CDR and by the University of Michigan Rackham Graduate Student Fellowship to ASV. The authors thank Osman Darici for helpful discussions concerning this work.

## Supporting Information

**S1 Text**

**Effects of Up and Down Steps on Rimless Wheel Step Timing and Average Forward Speed.**

## References

1. Davies S, Mackinnon, S. The energetics of walking on sand and grass at various speeds. Ergonomics. 2006;49(7):651–660.

2. Pandolf K, Haisman M, Goldman, R. Metabolic energy expenditure and terrain coefficients for walking on snow. Ergonomics. 1976;19(6):683–690.

3. Pinnington HC, Dawson, B. The energy cost of running on grass compared to soft dry beach sand. J Sci Med Sport. 2001;4(4):416–430.

4. Soule R, Goldman, R. Terrain coefficients for energy cost prediction. J Appl Physiol. 1972;32(5):706–708.

5. Voloshina AS, Kuo AD, Daley MA, Ferris, DP. Biomechanics and energetics of walking on uneven terrain. J Exp Biol. 2013;216(21):3963–3970.

6. Baratta R, Solomonow M, Zhou B, Letson D, Chuinard R, D’ambrosia R. Muscular coactivation: the role of the antagonist musculature in maintaining knee stability. The American journal of sports medicine. 1988;16(2):113–122.

7. Falconer K, Winter, D. Quantitative assessment of co-contraction at the ankle joint in walking. Electromyography and clinical neurophysiology. 1985;25(2–3):135–149.

8. Whitmore MW, Hargrove LJ, Perreault, EJ. Gait characteristics when walking on different slippery walkways. IEEE Transactions on Biomedical Engineering. 2016;63(1):228–239.

9. Blum Y, Birn-Jeffery A, Daley MA, Seyfarth, A. Does a crouched leg posture enhance running stability and robustness? Journal of Theoretical Biology. 2011;281(1):97–106.

10. Lejeune T, Willems P, Heglund, N. Mechanics and energetics of human locomotion on sand. J Exp Biol. 1998;201(13):2071.

11. Gates DH, Scott SJ, Wilken JM, Dingwell, JB. Frontal plane dynamic margins of stability in individuals with and without transtibial amputation walking on a loose rock surface. Gait & posture. 2013;38(4):570–575.

12. Gordon KE, Ferris DP, Kuo, AD. Metabolic and mechanical energy costs of reducing vertical center of mass movement during gait. Archives of physical medicine and rehabilitation. 2009;90(1):136–144.

13. Margaria R. Positive and negative work performances and their efficiencies in human locomotion. Int Z Angew Physiol. 1968;25(4):339–351.

14. Darici O, Temeltas H, Kuo, AD. Optimal regulation of bipedal walking speed despite an unexpected bump in the road. PLOS ONE. 2018;13(9):e0204205. doi:10.1371/journal.pone.0204205.

15. Kuo A. Energetics of actively powered locomotion using the simplest walking model. J Biomech Eng. 2002;124:113.

16. Kuo A, Donelan J, Ruina, A. Energetic consequences of walking like an inverted pendulum: step-to-step transitions. Exercise Sport Sci R. 2005;33(2):88–97.

17. McGeer T. Passive dynamic walking. Int J Robot Res. 1990b;9(2):62–82.

18. Alexander RM. Simple models of human movement. Applied Mechanics Reviews. 1995;48(8):461–470.

19. Garcia M, Chatterjee A, Ruina A, Coleman, M. The simplest walking model: stability, complexity, and scaling. J Biomech Eng. 1998;120(2):281–288.

20. Kuo A. A simple model of bipedal walking predicts the preferred speed-step length relationship. J Biomech Eng. 2001;123:264–269.

21. Mochon S, McMahon, TA. Ballistic walking. J Biomech. 1980;13(1):49–57.

22. Rebula JR, Kuo, AD. The cost of leg forces in bipedal locomotion: a simple optimization study. PloS one. 2015;10(2):e0117384.

23. Winter DA. Biomechanics and motor control of human movement. John Wiley & Sons; 2009.

24. Eng JJ, Winter, DA. Kinetic analysis of the lower limbs during walking: what information can be gained from a three-dimensional model? Journal of biomechanics. 1995;28(6):753–758.

25. Darici O, Temeltas H, Kuo, AD. Anticipatory control of momentum for bipedal walking on uneven terrain. Scientific Reports. 2019;Submitted, in review(bioRxiv 769828). doi:doi: https://doi.org/10.1101/769828.

26. Hof AL. Scaling gait data to body size. Gait & Posture. 1996;4(3):222–223.

27. Remy CD, Buffinton K, Siegwart, R. A matlab framework for efficient gait creation. In: 2011 IEEE/RSJ International Conference on Intelligent Robots and Systems. IEEE; 2011. p. 190–196.

28. Doke J, Kuo, AD. Energetic cost of producing cyclic muscle force, rather than work, to swing the human leg. J Exp Biol. 2007;210(13):2390–2398.

29. Maxwell Donelan J, Kram R, Arthur D K. Mechanical and metabolic determinants of the preferred step width in human walking. Proceedings of the Royal Society of London Series B: Biological Sciences. 2001;268(1480):1985–1992.

30. Huang PTW, Shorter KA, Adamczyk PG, Kuo, AD. Mechanical and energetic consequences of reduced ankle plantar-flexion in human walking. Journal of Experimental Biology. 2015;218(22):3541–3550.

31. Snaterse M, Ton R, Kuo AD, Donelan, JM. Distinct fast and slow processes contribute to the selection of preferred step frequency during human walking. Journal of Applied Physiology (Bethesda, Md: 1985). 2011;110(6):1682–1690. doi:10.1152/japplphysiol.00536.2010.

32. Su JLS, Dingwell, JB. Dynamic stability of passive dynamic walking on an irregular surface. J Biomech Eng. 2007;129(6):802–810.

33. Byl K, Tedrake, R. Metastable walking on stochastically rough terrain. Proc of Robotics: Science and Systems. 2008; p. 6490–6495.

34. Byl K, Tedrake, R. Metastable walking machines. Int J Robot Res. 2009;28(8):1040–1064.

35. Maus HM, Lipfert SW, Gross M, Rummel J, Seyfarth, A. Upright human gait did not provide a major mechanical challenge for our ancestors. Nature Comm. 2010;1:70.

36. Zelik KE, Kuo, AD. Human walking isn’t all hard work: evidence of soft tissue contributions to energy dissipation and return. The Journal of experimental biology. 2010;213(24):4257–4264.

